# T cell competition profoundly reduces the effect of initial precursor frequency on the generation of CD4 T cell memory

**DOI:** 10.1101/2020.09.09.290627

**Authors:** Adrian L. Smith, Barbara Fazekas de St. Groth

## Abstract

Protective immune responses are accompanied by increases in the frequency of high affinity T cells, which contribute to subsequent immunological memory. There is evidence that the fold-change in T cell number during the immune response is inversely related to initial precursor frequency, but the size of this effect remains poorly defined. Indeed, in many reports precursor frequency has been considered as directly proportional to the magnitude of the response. We have determined the effect of initial precursor frequency over the course of an in vivo antigen-specific response, in an experimental setting in which the other variables, TCR affinity and antigen dose, are kept constant. A major effect of precursor frequency was apparent in both the expansion and contraction phases; low initial precursor frequency in the physiological range was associated with greater initial expansion in T cell numbers, and also with preferential retention of memory cells. The effect was seen continuously across a 1000-fold naïve cell frequency range, leading to memory cell frequencies that differed by only 3-fold. These results are consistent with the existence of ongoing competition for antigen throughout the course of the immune response and explain the paradoxical ability of populations of genetically diverse individuals to make appropriate protective immune responses despite the large differences in initial repertoire that result from semi-random thymic TCR repertoire generation and selection.

## Introduction

Immunological memory gives rise to a more rapid and powerful response against an antigen previously encountered by the immune system. The state of immunological memory has been attributed to two independent mechanisms: an increase in the frequency of antigen-specific precursor cells, and changes in the behavior of individual lymphocytes, including altered expression of cell surface molecules involved in cell migration, adhesion, activation and cytokine signaling (1, 2). Such changes facilitate more rapid and effective activation of primed cells in comparison to naïve cells.

The frequency of antigen-specific T cells is known to increase acutely during primary immune responses. The response then enters a contraction phase in which many cells die, followed by a stable or memory phase. During this phase, cell numbers are still elevated over their initial levels, but the factors that determine memory cell numbers are still not fully established. Published reports contain widely differing estimates of the extent of both primary T cell clonal expansion and the size of the residual memory cell population.

Many factors may have contributed to these differing estimates, including detection methods with different sensitivities, analysis of different T cell subsets (3), and the extent of antigen persistence. Despite these sources of variation, one common trend has been for the initial precursor frequency of antigen specific T cells to be inversely related to the degree of expansion. Models in which the precursor frequency is high enough to be easily measured before immunization have yielded estimates as low as 2-10-fold for the increase in antigen specific cell number (4-7). In contrast, studies in which the initial precursor frequency was too small to be measured accurately have reported estimated increases of up to 2,000-fold for T cells expressing high affinity TCRs (8-10). One possible explanation for these findings is that T cells compete for access to antigen across a wide range of precursor frequencies, such that when precursors are abundant, proliferation is limited.

In addition to increases in precursor frequency, immunological memory is accompanied by an increase in the average affinity of the T cell receptors involved in the response. A number of studies have demonstrated that high affinity TCRs rapidly become dominant and are preferentially selected into the memory response (2, 11, 12). This is consistent with the existence of a mechanism of inter-clonal competition for antigen between T cells, such that cells of higher affinity proliferate and/or survive at the expense of cells with lower affinity TCRs.

We have previously described the effects of in vivo competition for antigen between T cells by examining the dynamics of T cell responses involving mixtures of T cells of defined affinity at unphysiologically high precursor frequencies (7). That study examined the simplest case of inter-clonal competition, namely competition for antigen between precursors of the same affinity. Competition limited T cell recruitment into division, burst size per recruited cell and, in the case of non-deletional responses, memory cell survival (7). To test whether competition for antigen was restricted to responses with high initial precursor frequencies, Quiel et al subsequently performed a similar experiment at lower precursor frequencies, measuring T cell proliferation over the first 7 days of the in vivo response for a 100-fold range of input cell doses (13). Once again, a competitive effect was observed. The data from this limited timeframe were consistent with multiple models that postulated different mechanisms to explain quantitative aspects of the response (14, 15).

Here we report the effect of a 1000-fold range of precursor frequencies on the response of TCR transgenic T cells to a defined dose of antigen, extending the analysis to include the 6-week timepoint, which is of crucial importance in defining the memory response. We transferred TCR transgenic T cells into groups of unmanipulated adoptive hosts and used flow cytometry to track responses to subcutaneous immunization with a defined dose of specific peptide/adjuvant. This enabled us to estimate when competition was occurring, and whether it was continuous over the entire range of precursor frequencies, including those in the physiological range. By the peak of the response on day 7, the 1000-fold difference in antigen-specific cell number between the groups had been reduced to 12-fold. At day 42, by which time significant contraction in Ag-specific cell numbers had occurred, the difference was only 3-fold. These results strongly suggest that competition occurs over a wide range of precursor frequencies, both during the expansion and contraction phases of immune responses, such that the final effect of vastly different initial precursor frequencies on both the primary and secondary immune responses is profoundly reduced.

## Results

In previous studies, we compared Ag-specific CD4^+^ T cell responses in animals in which Tg T cells made up 90% or 2% of peripheral CD4 T cells (7). Competition between T cells at the higher precursor frequencies was indicated by a reduction in the proportion of precursors recruited into the response and in the burst size per recruited precursor. The competition operated principally at the level of specific peptide-MHC complexes rather than antigen-nonspecific APC factors, and dose-response curves indicated that antigen dose rather than initial precursor frequency was the major driver regulating the size of the immune response. Moreover, precursor frequency did not change the character of the response, in that intravenous peptide induced deletion of responding cells, whereas subcutaneous peptide plus adjuvant induced priming, regardless of precursor frequency.

A significant limitation of our earlier study was the necessity of using high precursor frequencies so that the events following initial activation could be easily followed by flow cytometry. In order to extend the study to lower, more physiological, cell numbers, we transferred 10^4^, 10^5^, 10^6^ or 10^7^ Tg LN cells labeled with 5-(and-6)-carboxyfluorescein diacetate succinimidyl ester (CFSE) into non-Tg CD45.1 congenic recipients. Four days after transfer, 3 mice from each group were sacrificed and LNs and spleen examined by flow cytometry to ascertain the efficiency of adoptive transfer and establish the initial number of precursors present in each group. Three days later, the remaining 8 mice per group were immunized subcutaneously with 10ug of moth cytochrome c (MCC_87-103_) peptide emulsified in Complete Freund’s Adjuvant (CFA). Draining LNs, distal LNs, spleens and peripheral blood were examined by flow cytometry on days 7 and 42 after immunization, representing points near the peak of the response and late in the contraction phase, respectively.

Figure 1 shows the kinetics of the response in the draining LNs, expressed as the absolute number of transgenic (CD45.1-negative) T cells (Fig 1a) or as the number of transgenic T cells per 10^5^ leukocytes (Fig 1b). Comparison of the data derived from the 2 different analyses indicated that the increase in transgenic cells per lymph node on day 7 was larger than the increase per 10^5^ leukocytes, suggesting that there had been an influx of antigen non-specific leukocytes into the draining LN during the expansion phase of the response. The most striking feature of the data was the dramatic narrowing of the range of responder cell numbers as the response progressed. Log transformation of the input and output cell numbers for each animal revealed a clear linear relationship at each time point, with the slope decreasing over the course of the response (Figs 1c,d), indicating that the output numbers were becoming progressively closer to each other. The fact that the log/log plots remained linear indicates that competition for antigen occurs as a continuous process at all input cell numbers, rather than a discontinuous process that affects only high precursor numbers, as suggested by others (16). Moreover, the effect could be clearly seen in both the expansion and contraction phases of the response, suggesting that competition between T cells is a fundamental and continuous property of the immune response.

**Figure 1:**
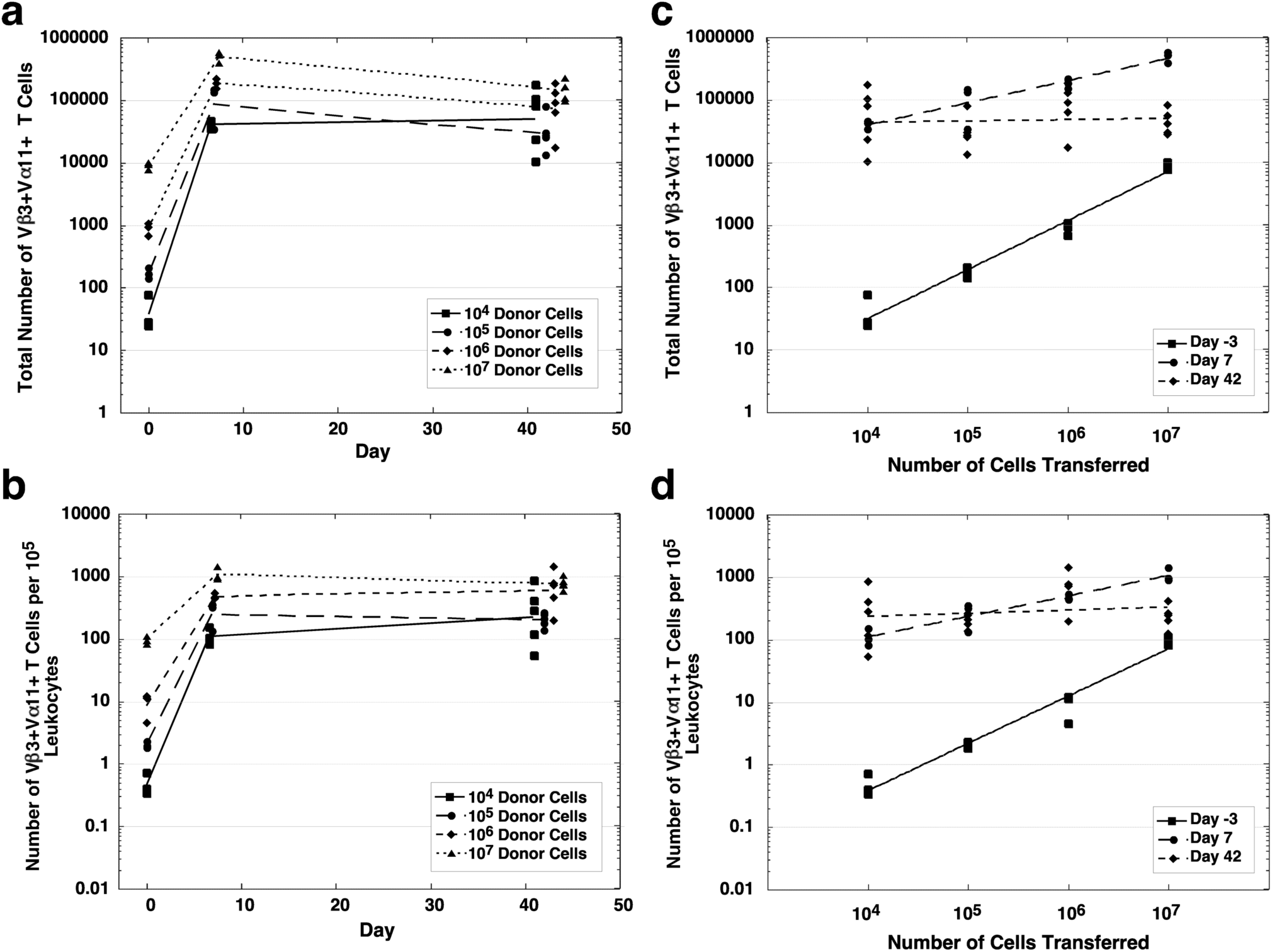
Relationship between initial precursor frequency and the size of the expansion and contraction phases in response to subcutaneous immunization. Kinetics of the response in the draining LNs to s.c. peptide in CFA in adoptive non-Tg hosts of 10^4^, 10^5^, 10^6^ or 10^7^, lymphocytes from -D TCR Tg mice. Draining LN were analysed three days before immunization (shown as time point 0 in the graph) and on days 7 and 42 after immunization. Symbols represent individual mice (3-5 per group, per time point). a) Total number of CD4 T cells expressing the Tg TCR. Numbers were calculated by multiplying the percentage of cells determined by flow cytometry by the total number of leukocytes recovered from each organ. b) Number of CD4 T cells expressing the Tg TCR per 10^5^ total live leukocytes as defined by forward/side scatter and exclusion of PI. c) Relationship of total number of transferred CD4 T cells expressing the Tg TCR to the number recovered from draining LNs. The line of best fit for each time point is shown. d) Relationship of total number of transferred CD4 T cells expressing the Tg TCR per 10^5^ total live leukocytes to the number recovered from draining LNs. The line of best fit for each time point is shown.

In contrast to our previous study in which CFSE division profiles indicated an inverse relationship between precursor frequency and burst size, all the responding cells in this study had divided to become CFSE negative by day 7. Representative dot plots for each cell dose and time point, gated for donor-derived (CD45.1^-^) CD4 cells, are shown in Figure 2. In addition to divided transgenic T cells (CFSE-negative and Tgα^+^), these plots show a variable population of Tgα^-^ CFSE-negative cells, even in unimmunized (day –3) mice. Analysis of this population in unimmunized and PBS-immunized animals revealed that these cells were small resting cells that stained positively for CD4 but were predominantly CD44^low^ (data not shown). These characteristics suggested strongly that they were not divided donor-derived cells, but rather host-derived cells that had failed to stain with anti-CD45.1. Because a small population of similar cells also contaminated the Tgα^+^ gate used to calculate the number of antigen-specific cells on days 7 and 42 (boxed in Fig. 2), these cells had a minor effect the analysis of the experiment. To offset this technical artifact, we devised a correction based on the assumption that the ratio of Vα11^+^ to Vα11^-^ cells within the contaminant population was constant. This assumption was confirmed by demonstrating the linearity of plots of the number of Vα11^-^ versus Vα11^+^ (Tgα^+^) cells within the CFSE negative region in unimmunized or PBS/CFA immunized animals on day 7 (Fig. 3a) and day 42 (Fig. 3c). These plots indicated that the number of host CFSE^-^ cells contaminating the Vα11^+^ gate was directly proportional to the number of CFSE^-^ Vα11^-^ cells (R^2^ between 0.958 and 0.999). Using the equations generated from these plots, corrected numbers of Tgα^+^ cells were calculated for days 7 and 42 (Figs 3b and 3d, respectively). As expected, these corrections had the greatest effect on the numbers calculated from mice that received the lower doses of cells, and at the last time point, when the contaminating population contributed a greater proportion of the total event count.

**Figure 2:**
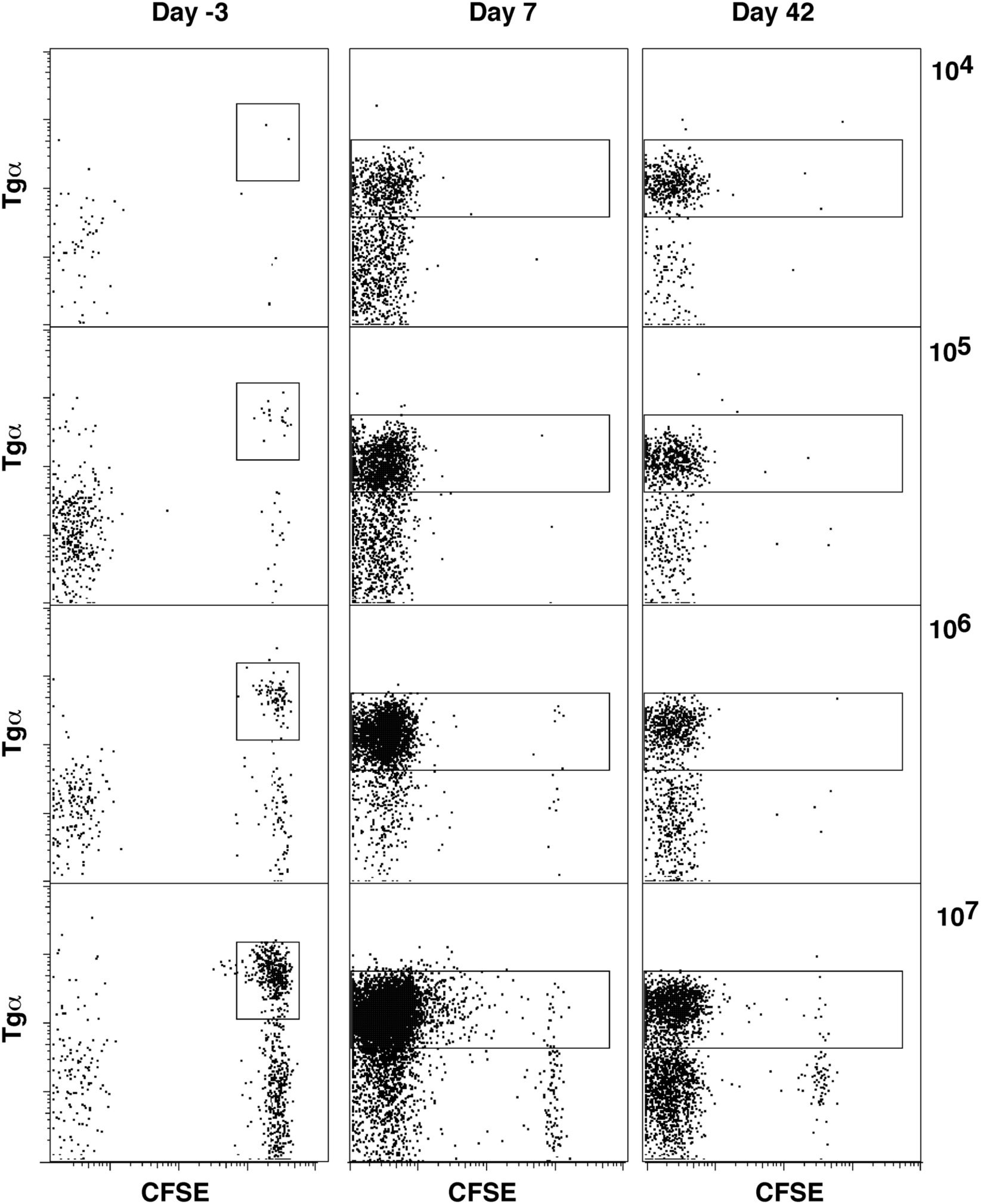
CFSE profiles of CD45.1^-^ CD4 T cells in draining LNs. Representative dot plots for each cell dose and time point. Cells are gated for forward/side scatter, lack of staining for CD8, Mac1, B220 and CD45.1, expression of CD4 and Vβ3 and exclusion of PI. Boxes enclose cells expressing the Tgα chain (Vα11).

**Figure 3:**
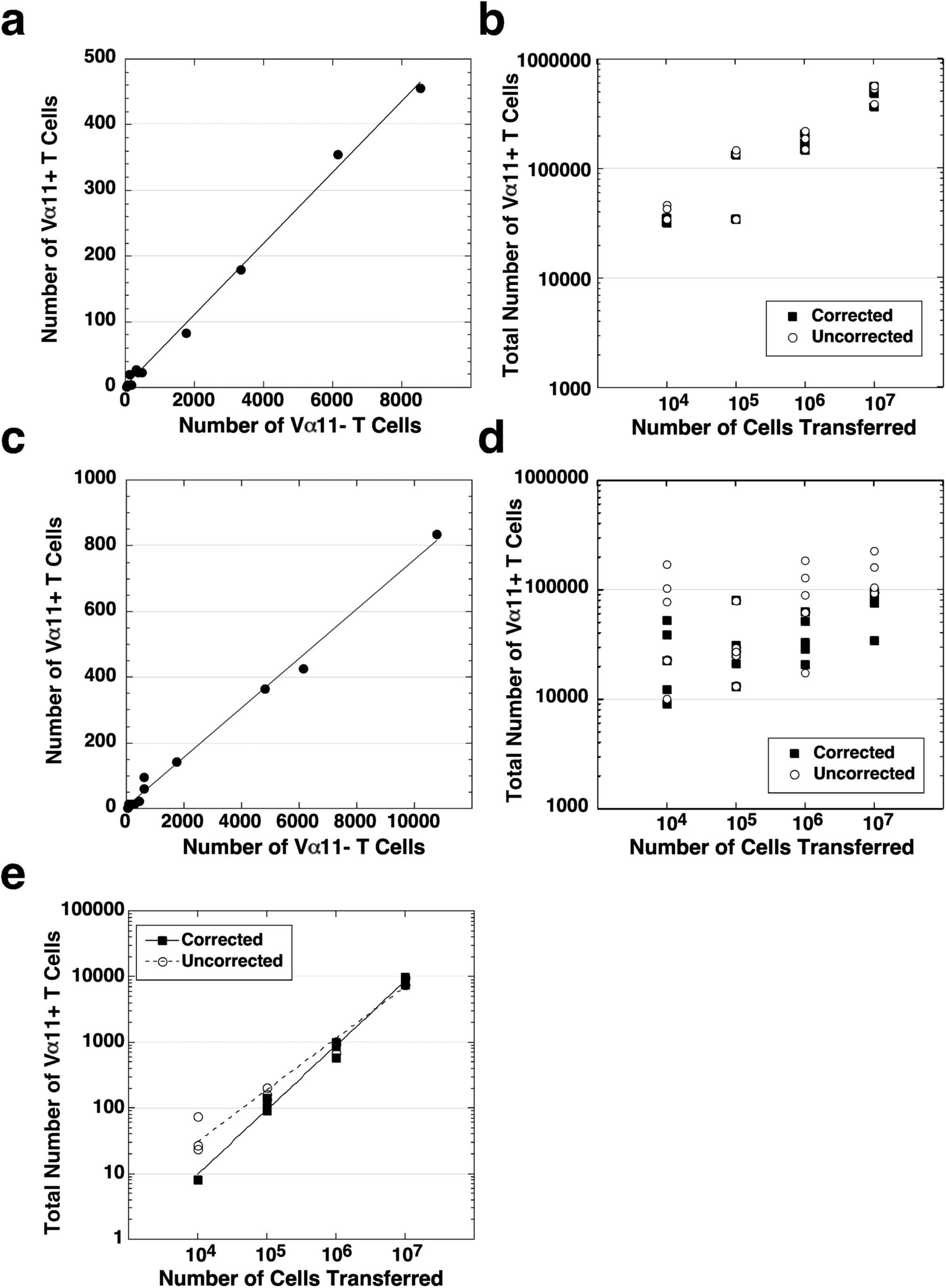
Corrections for confounding cell populations and cytometer background. a) Analysis of day 7 LN from unimmunized and PBS/CFA immunized mice. Plots show the numbers of Vα11^+^ vs Vα11^-^ cells within CFSE-negative CD4^+^Vβ11^+^ cells (gated as negative for CD8, Mac1, B220, CD45.1 and PI). These cells appear to be of host-origin as they are small, resting and CFSE-negative, while donor cells are uniformly undivided CFSE^+^. The line of best fit is indicated (R^2^ = 0.999). b) Number of Tgα^+^ CD4 T cells in the draining lymph nodes on day 7, with corrected numbers after removal of Vα11^+^ cells calculated using the line of best fit from a). Similar calculations were carried out for the distal LNs and spleen (not shown). c) Analysis of day 42 LN from PBS/CFA immunized mice. Plots show the numbers of Vα11^+^ vs Vα11^-^ cells within CFSE-negative CD4^+^Vβ11^+^ cells (gated as negative for CD8, Mac1, B220, CD45.1 and PI). The line of best fit is indicated (R^2^ = 0.958). d) Number of Tgα^+^ CD4 T cells in the draining LN on day 42 with corrected numbers after removal of Vα11^+^ cells calculated using the line of best fit from c). e) Number of Vα11^+^ CD4 T cells in the lymph nodes on day −3. Corrected numbers were calculated by removal of the number of cells due to cytometer background. Background was determined by running identically stained samples without a CFSE labeled population. This reduced the events in some samples of the 10^4^ group to zero, ie below the limit of detection. Similar calculations were performed for spleen (not shown).

Fig. 2 also shows that before immunization (day –3), the number of events falling within the CFSE^+^Tgα^+^ gate was very small at the lowest dose of donor cells. At this level, background events from electronic noise or debris within the fluid lines of the flow cytometer can become a significant contribution to the total event count. The level of this background was calculated by running identically stained samples from mice that had not received a CFSE-labeled cohort of cells (data not shown). The corrected cell numbers are plotted in Figure 3e, showing that once the background is removed, the expected linear relationship between the cell doses is restored. The removal of the background for the 10^4^ input cell group reduced the event count for 2 out of 3 mice to zero, indicating that the limit of detection by flow cytometry was being reached at this number of donor cells.

The corrected kinetics for draining lymph nodes, distal lymph nodes and spleen are shown in Figs 4a-c. The overall pattern of kinetics is the same as in the uncorrected plots (Fig. 1), indicating that the process of correction did not change the overall conclusions of the study, but simply reduced the level of “noise” in the data. For example, the curves for the different cell doses no longer cross each other (compare Figs 1b and 4a), and the log/log plots for each dose at each time point show a slightly smaller range of cell numbers. The number of cells in the spleen on day –3 after inoculation with 10^4^ donor cells was below the limit of detection in all three mice in the group.

**Figure 4:**
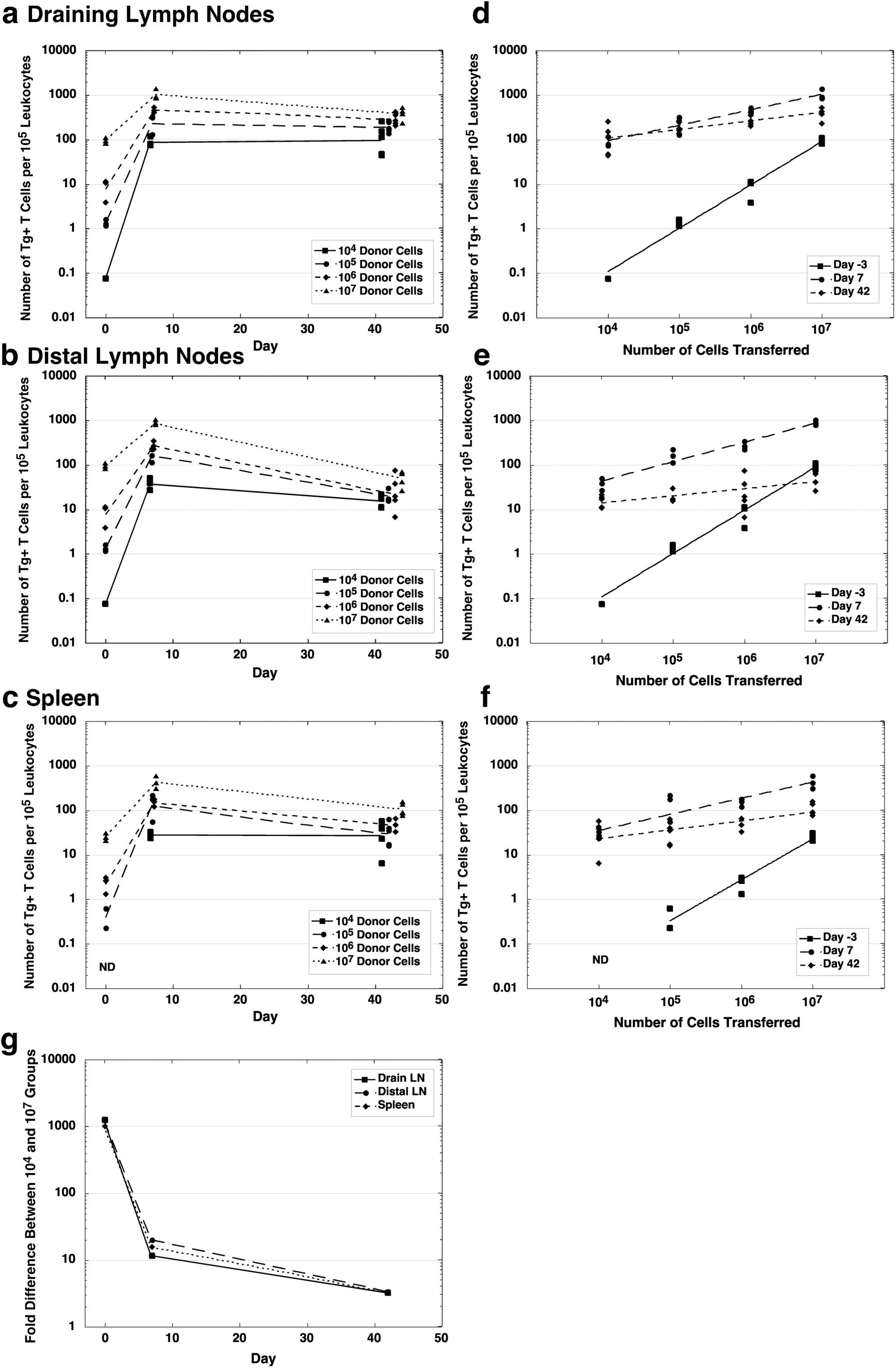
Corrected kinetics of the response to MCC_87-103_/CFA. Cell numbers calculated per 10^5^ leukocytes as for Figure 1b with corrections described in Figure 3. a) Corrected kinetics in draining LNs. b) Corrected kinetics in distal LNs. c) Corrected kinetics in spleen. Subtraction of the background on day −3 indicated that the number of cells recovered from the 10^4^ group was below the limit of detection. d-f) Relationship between the number of cells transferred to the number recovered from draining LNs (d), distal LNs (e), spleen (f). The line of best fit for each time point is shown. g) Fold difference between the mean number of Tg^+^ CD4 T cells per 10^5^ leukocytes in the 10^4^ and 10^7^ groups at each time point.

As described above, the slope of the line of best fit for the log/log plots (Figs 4d-f) decreases with each time point, demonstrating that the cell numbers in different groups are becoming closer throughout the response. This is summarized in Figure 4g, which shows that an initial 1000-fold difference on day –3 had decreased to approximately 12-fold by day 7 and further narrowed to less than 3-fold by day 42.

A similar level of expansion in cell numbers at day 7 was seen in both the draining and distal LNs (Figs 4a-b), although initial proliferation is clearly restricted to the draining nodes in this model, suggesting that cells were rapidly being exported from the draining node during the early phases of the immune response (17). However, the cell numbers contracted less between d7 and d42 in the draining than the distal LNs. This is shown most clearly by plotting the number of cells in the draining versus the distal lymph nodes (Figure 5a). This plot shows that at d7 the cell numbers in the two sites retained the same relationship they had before immunization, while by d42 there were relatively more cells in the draining node. In contrast, the draining LN and spleen retained the same relationship at all three time points (Figure 5b). The straight-line relationships for the day −3 and day 7 data in these two plots suggest that export from the draining LN before day 7 was directly proportional to cell number. Between d7 and d42, the change in numbers in LNs versus spleen are consistent with restricted entry to LNs but not spleen due to the loss of CD62L expression by effector memory T cells (18). The relative preservation of cell number in the draining LNs may have been due to antigen persistence in those LNs or the tissues they drain. This would serve to keep the cell numbers elevated despite loss of CD62L, either by limiting export from the nodes, slowing death, keeping some cells in cycle or, most likely, by recirculation of effector/memory cells through the peripheral tissue (footpad) and back into the node via the afferent lymph rather than via the blood and high endothelial venules. Persistence of antigen is supported by the observation that Tg cells in the draining LN were on average larger than the Tg cells in the distal LN (forward scatter of 145 in draining LN versus 128 and 130 in distal LNs and spleen, respectively) and had a higher level of CD69 expression (Figures 5c and 5d), both consistent with recent antigen exposure.

**Figure 5:**
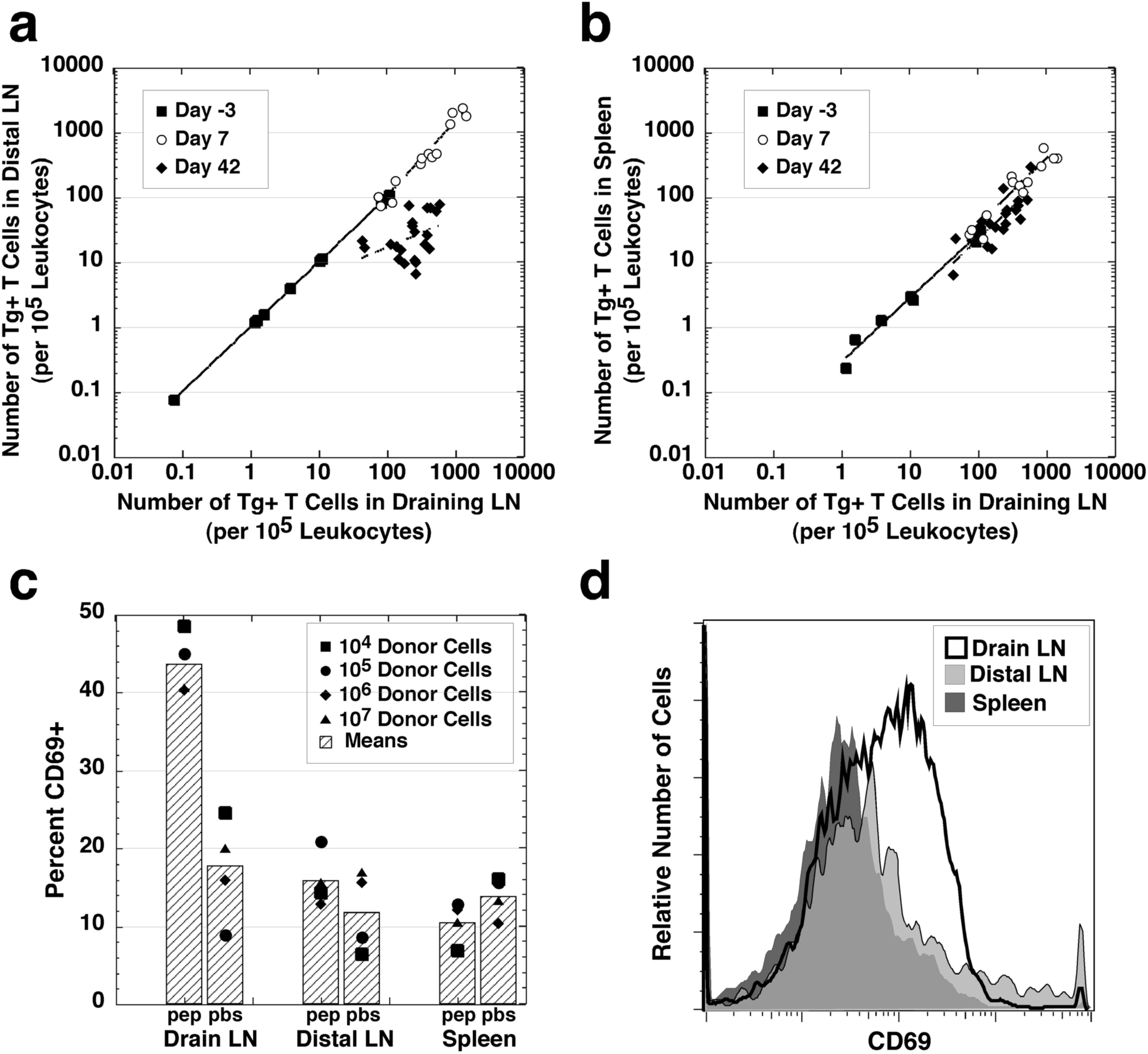
Comparison of cells in the draining LNs, distal LNs and spleen. a) Number of Tg^+^ CD4 T cells (per 10^5^ total leukocytes) in the draining LNs versus the distal LNs. Three time points shown each with line of best fit. Cell numbers calculated as in Figure 1b with corrections applied. b) Number of Tg^+^ CD4 T cells (per 10^5^ total leukocytes) in the draining LNs versus the spleen. Three time points shown each with line of best fit for each. Cell numbers calculated as in Figure 1b with corrections applied. c) Percentage of Tg^+^ CD4 T cells expressing CD69 on day 42 in mice immunized with MCC_87-103_/CFA (pep) or PBS/CFA (pbs). Cells gated for forward/side scatter, lack of expression of Ly5.1, expression of CD4 and Vα11^+^, and exclusion of PI. d) Histogram of CD69 expression on day 42 in draining and distal LNs and spleen of MCC_87-103_/CFA immunized mice. Cells gated as in c).

## Discussion

Studies of in vivo T cell responses have produced widely varying estimates of the extent of clonal expansion and contraction. Early studies of the response to infectious organisms suggested a 1000-5000 fold expansion, followed by a 10-100 fold contraction (19-22), whereas studies using TCR Tg mice showed only 10-fold expansion with very little evidence of a prolonged increase in precursor frequency (4-6). One study of an individual high affinity endogenous clone, detected by specific TCR CDR3 sequences, suggested up to 2,000-fold expansion (2). Several possible reasons have been advanced to explain these differences, including inherent differences between CD4 and CD8 T cells, differences in antigen (live versus model peptide), different methods of quantifying memory T cells, differences in antigen persistence and even artifacts due to transgenic systems (23).

The early studies demonstrating competition between T cells for Ag-MHC complexes (7, 24) suggested an alternative explanation to account for the observed differences in T cell burst size in different models. If T cells compete for antigen, then in a given system in which antigen is limiting, a larger initial precursor number will result in a smaller estimate of clonal expansion. This is consistent with the observation that those studies in which the precursor frequency was too small to be measured accurately yielded estimates of fold expansion at the upper end of the range while in those with higher initial frequencies (i.e. in the range that could be directly measured before immunization), the reported fold increase was far lower.

To directly address the question of whether competition occurs at precursor frequencies in the physiological range, we adoptively transferred CD4 Tg T cells over a 1000-fold range of cell numbers and followed their response to subcutaneous antigen in adjuvant. We were able to detect cells with an initial precursor frequency as low as 0.00017% of CD4^+^ T cells (or 1.7 cells per 10^6^ CD4 cells), a level consistent with estimates of precursor frequency in normal naïve mice (24-28). By following the Tg T cell response to subcutaneous immunization, we have shown that competition occurs over a wide range of precursor frequencies and affects both the expansion and contraction phases of the response. The initial 1000-fold difference in specific cell numbers was reduced to 12 fold at the peak and to less than 3 fold by day 42, providing a striking demonstration of how the immune system compensates for large differences in the initial precursor pool. Moreover, the fact that competition is an intrinsic part of the immune response was apparent from the clearly continuous function governing log input to output cell number over a wide range of precursor frequencies (Fig 4d-f).

Multiple reports have suggested or assumed that the size of T cell response is determined by initial naive precursor cell frequency (28-33). In many of these studies, precursor frequency, TCR affinity and efficiency of epitope presentation were all variable, so that the effect of precursor frequency alone could not be determined. In addition, while a strong correlation clearly exists between precursor frequency and burst size, our quantitative studies indicate that burst size increases only 1.12-fold for every 10-fold increase in initial precursor frequency, when TCR affinity and initial antigen dose are kept constant.

Interestingly antigen-specific cell numbers appeared to contract more in the distal LNs than the draining LN and the spleen. Two processes probably underlie this finding. First, activated /memory cells would be expected to be at least partially excluded from the peripheral lymph nodes compared to the spleen, due to changes in expression of homing molecules such as CD62L (34, 35), explaining why the distal nodes have a lower frequency of cells than the spleen. However draining nodes would also receive additional cells draining from the depot of persistent antigen at the site of injection, via the lymphatics to the node. The results presented here suggest that antigen persistence may be an additional factor contributing to differences in the degree of contraction observed in different models. For example, LCMV in mice (36) and EBV in humans (37) have been shown to persist in lymphoid tissues for relatively long periods, perhaps partly explaining why memory cells remain at much higher levels after these infections (8, 21, 24, 38) than after other viral responses (20, 39).

In our previous study (7), the number of T cells persisting at six weeks was only between 2 and 4-fold higher than the initial frequency, whereas in this study the increase was 10-fold in the group with the most precursors and ∼1000-fold in the group with the least. This difference can be explained by the higher initial precursor frequency in the previous study (2% of peripheral CD4 T cells versus 0.2% in the current study). Thus in different studies using the same system we have demonstrated that changes in initial precursor frequency profoundly influence the calculation of the fold increase in T cell numbers during a response, but have only a limited impact on the actual number of memory T cells generated by the response. Č

Indirect evidence of interclonal competition between T cells of different affinities has previously been derived from studies of average T cell affinity in primary and secondary immune responses (12, 25) and from observations that the average TCR affinity is an inverse function of Ag dose in vivo (40, 41). By demonstrating that competition can occur even at extremely low numbers of high affinity cells, our study demonstrates a mechanism that would be expected to play an important role in the affinity maturation of a T cell response by ensuring that the few naïve cells of the highest affinity can expand sufficiently to dominate the response. The small number of very high affinity cells Tg^+^ cells introduced by adoptive transfer into naïve hosts with an endogenous CD4 T cell repertoire were able to proliferate to the extent that they made up approximately 0.5% of peripheral CD4 T cells, a level consistent with the total specific response in previous studies of the polyclonal response to cytochrome *c* (2), and in CD4 T cell responses to viruses (42).

Indirect studies of affinity maturation of the T cell response have led to differing conclusions regarding the timing of the process. Some indicate that repertoire narrowing can occur early in the primary response (8, 25, 27, 43, 44), others did not see narrowing until the contraction phase (45), and yet others found no evidence of selection until the secondary response was underway (12, 46). While the basis for these differences remains unclear, the current study, which utilizes a model peptide antigen, clearly demonstrates that competition can occur both during the expansion and contraction phases of the primary response. When the antigen is a proliferating virus or bacterium, it is possible that by the time the immune system recognizes the threat, enough antigen may be present to minimize the effect of competition in the primary response, particularly in the case of systemic infections. However the more rapid memory response to secondary challenge will eliminate the organism more quickly, limiting the amount of antigen and resulting in more easily detectable competition between T cells.

The mechanism of T cell interclonal competition for antigen is unknown. In contrast to antigen sequestration via internalization by B cells, there is only very limited data regarding down-modulation of specific peptide-MHC complexes during the in vivo T cell response. Kedl et al used a monoclonal antibody specific for an OVA-K^b^ complex to demonstrate downregulation during a CD8 T cell response (47). Unfortunately a high affinity, highly specific monoclonal recognising MCC-IE^k^ complexes was not available for our study, and attempts to measure antigen levels by performing ex vivo T cell activation studies in response to DCs purified at different stages of the in vivo immune response were not successful, due to the relative antigen insensitivity of in vitro proliferation responses. It may be possible to measure specific MCC-IE^k^ complexes expressed by ex vivo DCs using a direct visualisation technique such as those developed by the Davis lab to image very early T cell activation events at the single cell level (48, 49).

In summary, this study demonstrates that T cell competition occurs at physiological precursor numbers, during both the expansion and contraction phases of a T cell response. It provides evidence supporting a model in which competition for Ag allows the immune system of outbred species to make appropriate responses to both self and foreign antigens, regardless of the wide variations in affinities and frequencies of the TCRs selected by the combination of MHC-antigen complexes present in each individual. Competition for antigen would not only ensure that clones of the highest affinity are preferentially expanded in the primary response and selected into the memory population, but would also to allow the amount of available antigen to serve as the principal determinant of the final size of the T cell response.

## Methods

### Experimental Animals

Transgenic (Tg) mouse lines were bred and housed under specific pathogen-free conditions at the Centenary Institute Animal Facility. Approval for all animal experimentation was obtained from the Institutional Ethics Committee at the University of Sydney. The -D line TCR Tg line expressing the 5C.C7 receptor, which recognizes the COOH-terminal epitope of MCC in the context of IE^k^ (50, 51) was maintained on a B10.BR (H-2^k^) background. For the experiments in this study, the -D line was crossed with C57BL/6 to provide F1 (H-2^bk^) animals. Recipient mice were F1 (H-2^bk^) crosses of B10.BR with the C57BL/6 Ly5.1 congenic line B6.SJLPtrpc^a^ (Animal Resources Centre, Perth, Australia).

### T Cell Purification, Labeling, and Injection

Pooled inguinal, axillary, subscapular, cervical, and para-aortic LNs of naive F1 (5C.C7 TCR x C57BL/6) mice served as the source of MCC-specific T cells, which were labeled with 5-(and-6)-carboxyfluorescein diacetate succinimidyl ester (CFSE; Molecular Probes, Eugene, OR) as described (52). Each recipient received between 10^4^ and 10^7^ labeled cells (containing approximately 30% MCC-specific CD4^+^ T cells) intravenously via the lateral tail vein.

### Antigens and Immunization

MCC peptide 87-103 was purchased from Chiron Technologies (Melbourne, Australia). For s.c. immunization, peptide was dissolved in PBS, emulsified 1:1 in Complete Freund’s Adjuvant (Commonwealth Serum Laboratories, Melbourne, Australia) and a total volume of 200ul was distributed between injection sites in both hind footpads (50ul each) and base of tail (100ul). Control mice received 100ul PBS emulsified in 100ul CFA.

### Flow Cytometry

Five colour antibody staining was performed in 96-well round-bottom microtitre plates (ICN, Costa Meas, CA) as described previously (17, 53). CFSE fluorescence of transferred T cells was detected in the FL-1 channel. For determining T cell numbers the following staining strategy was used: CD4 was detected in FL2 using anti-CD4-PE (RM4-5; PharMingen, San Diego, CA); FL3/1 was used as a dump channel to exclude both dead cells (propidium iodide (PI) positive) and cells that were not of interest by applying a cocktail of biotinylated antibodies, consisting of anti-CD8α (53-6.7; PharMingen), anti-Mac1 (M1/70;PharMingen), anti-B220 (RA3-3A1, conjugated in-house) and anti-CD45.1 (A20.1, conjugated in-house), plus streptavidin-QuantumRed (Sigma, St Louis, MO). Tgβ chain was detected in FL4 using anti-Vβ3 supernatant (KJ25) plus anti-hamster TexasRed (Caltag, Burlingame, CA); Tgα chain in FL5 was detected with anti-Vα11 supernatant (RR8.1) plus anti-mouse-allophycocyanin(APC) (rat Ig cross-reactive; Biomeda, Foster City, CA). For detecting CD69 expression on CD4 T cells, anti-CD4-PE was combined with anti-CD69bi (conjugated in-house) detected with streptavidin-Alexa Fluor™ 594 (Molecular Probes) and anti-CD45.1-APC (A20.1, conjugated in-house).

7-channel data acquisition (0.5–1 × 10^6^ events per sample) was performed on a FACStarPLUS^®^ flow cytometer (Becton Dickinson, Mountain View, CA) equipped with a 488-nm argon-ion laser and a 610-nm dye laser and analyzed using FlowJo software (Tree Star, San Carlos, CA) following the gating strategy outlined previously (54).

## Abbreviations used in this paper

CFA: Complete Freund’s adjuvant
CFSE: 5-(and-6)-carboxyfluorescein diacetate, succinimidyl ester
MCC_87-103_: moth cytochrome *c*, peptide residues 87-103
PI: propidium iodide
Tg: transgenic.

## Acknowledgments

The authors wish to thank Karen Knight and the staff of the Centenary Institute Animal Facility for excellent animal husbandry, and Kate Scott and Sarah Smith for managing the Tg colony and screening the mice used in this study. Barbara Fazekas de St. Groth was supported by a Wellcome Trust Senior Research Fellowship and a National Health and Medical Research Council of Australia Principal Research Fellowship, and Adrian Smith by an Australian Postgraduate Award. This work was funded by the National Health and Medical Research Council of Australia and the Wellcome Trust. The support of the New South Wales Health Department through its research and development infrastructure development grants program is gratefully acknowledged.

